# Comparison of the effects of lithium orotate and lithium carbonate on locomotion and memory in a *Drosophila melanogaster* model of Alzheimer’s disease

**DOI:** 10.1101/2025.08.13.670149

**Authors:** Ishan Reddy Alla, Ryan Chung

## Abstract

Alzheimer’s disease (AD) is a terminal neurodegenerative disease characterized by cognitive decline and memory loss resulting from the buildup of amyloid-beta plaques and tau protein tangles. Lithium salts, particularly lithium carbonate (Li_2_CO_3_), inhibit the enzyme Glycogen Synthase Kinase-3, which is upregulated in AD, reducing tau and amyloid-beta accumulation, inflammation, and oxidative stress. However, Li_2_CO_3_ has declined in use due to long-term neurotoxicity. Lithium orotate (LiOr), an alternative lithium salt, has greater bioavailability–delivering more ions to the brain–but was discontinued in the 1970s due to toxicity concerns at equivalent doses. It was hypothesized that LiOr would be more effective than Li_2_CO_3_ at lower doses (5 mM) due to higher lithium ion transfer, while Li_2_CO_3_ may outperform LiOr at higher doses (10 mM) due to LiOr’s toxicity. This study compares LiOr and Li_2_CO_3_ in treating locomotion and memory in a *Drosophila melanogaster* model produced by crossing UAS amyloid-beta 42 and Act5c GAL4 flies via the GAL4/UAS system. Once aged to 25 days, flies were assessed for locomotion using the negative geotaxis assay and for short-term memory using the aversive phototaxic suppression (APS) assay. These behavioral assays aim to quantify changes in cognitive and motor function resulting from treatment. Results suggest Li_2_CO_3_ improves cognitive and motor performance in healthy flies, but LiOr was not found to be significantly more effective. This study offered a novel comparison of two lithium compounds in an Alzheimer’s model and may guide research into safer, more effective AD and lithium treatments.

## Introduction

Alzheimer’s Disease (AD) is a terminal neurodegenerative disease, the most common form of dementia, that affects an estimated 6.7 million seniors living in the United States, a number expected to almost triple by the year 2060 (“Alzheimer’s & Dementia”, 2023; Rocca et al., 2011). Characterized by progressive memory loss, cognitive dysfunction, and neuronal death, AD is pathologically marked by the accumulation of amyloid-beta plaques, tau protein tangles, and synapse loss (Tue et al., 2020) linked to upregulated enzymes such as Glycogen Synthase Kinase-3 (GSK-3) which disrupts key cellular processes (Hooper et al., 2008). AD’s upregulation of these enzymes exacerbates neuroinflammation (immune response to damage in neurons) and mitochondrial dysfunction (Alzheimer’s Association, n. d.).

Of the several hypotheses proposed to explain AD pathology, the amyloid-beta (Aβ) hypothesis is the best-developed, suggesting that an imbalance in the production and clearance of Aβ peptides forms Aβ plaques that disrupt brain function and trigger the remainder of AD processes (Hardy, 2002).

The immediate cause for AD onset remains unknown; a combination of genetic, environmental, and lifestyle factors are found to contribute to disease onset (“What causes Alzheimer’s disease”, 2024). As of now, AD has no cure, however, several treatments aim to improve the quality of life for patients by alleviating symptoms. Despite eight FDA-approved AD medications, only two target disease progression (via reducing Aβ levels) (National Institute on Aging, 2024), but even those have limited success in improving cognitive function and are often accompanied by severe side effects such as brain swelling, bleeding, nausea, and dizziness (Shen et al., 2024). This lack of an effective treatment underscores the need for further pharmacological research to develop safer and more effective treatments.

Lithium, historically successful in mood stabilization, has shown promise for its neuroprotective potential in neurodegenerative diseases such as AD (Brown & Tracy, 2013).

Lithium’s inhibition of GSK-3 prevents β-catenin degradation (allowing β-catenin to accumulate and regulate gene transcription in the nucleus), which ultimately reduces Aβ deposition, enhances neuron growth, and improves cognitive function in AD (Shen et al., 2024). Numerous animal studies have demonstrated that lithium treatment reduces Aβ and tau accumulation, attenuates neuronal cell death, and improves cognitive function. Shin et al. (2023) concluded that most studies have consistently reported that lithium reduces Aβ aggregation, phosphorylated tau formation, and neuronal death and improves learning and working memory in APP mouse models. Sofola-Adesakin et al. (2015) determined that, in an AD *Drosophila* model, lithium caused a reduction in Aβ42 protein synthesis. At all doses tested in the experiment, lithium attenuated the locomotory defects induced by Aβ42, but it rescued lifespan only at lower doses, suggesting that long-term, high-dose lithium treatment may have induced toxicity.

The most effective lithium drug in disease treatment, Li_2_CO_3,_ has a narrow therapeutic index (range of dosages with effective treatment) along with side effects ranging from inconvenient, such as nausea, to life-threatening, such as renal dysfunction (Pacholko & Bekar, 2021). Research on Li_2_CO_3_’s neurotoxicity after extended usage has caused a rapid decline in usage over two decades (Pérez de Mendiola et al., 2021).

Another lithium drug, lithium orotate (LiC_5_H_3_N_2_O_4_; referred to as LiOr), is an aqueous lithium salt of lithium and orotic acid. Because orotic acid is a bioavailable (effective ion transmitter) mineral carrier that can more readily transport inorganic ions across biological membranes with a more consistent Li^+^ ion transfer than Li_2_CO_3_, LiOr is predicted to cross the blood–brain barrier and enter cells more readily than Li_2_CO_3_, reducing dosage requirements and lessening toxicity concerns for LiOr (Pacholko & Bekar, 2021).

After LiOr was introduced, studies using rodent models determined that LiOr was more bioavailable than Li_2_CO_3_. In 1973, Hans Nieper found that LiOr yielded higher brain concentrations of lithium than Li_2_CO_3_, but possibly had higher toxicity. In 1978, a research experiment found that equivalent dosages of LiOr and Li_2_CO_3_ led to LiOr having 3x the lithium transfer into the brain and a much more progressive pattern of lithium transfer compared to Li_2_CO_3_ (Kling et al.).

Although evidence for enhanced brain bioavailability in LiOr was initially found, research into LiOr was discontinued for decades due to renal toxicity concerns with equivalent doses of LiOr and Li_2_CO_3_ (Smith & Schou, 1979). These experiments failed to test for LiOr’s effectiveness at lower doses than Li_2_CO_3_ due to its bioavailability, and as such, currently, there is a notable gap in the data regarding LiOr’s therapeutic index, particularly its impact on behavioral symptoms. LiOr has never been tested to see how varying doses affect any behavioral symptoms in AD. In the past decade, reviews have highlighted the wealth of shortcomings in these past comparisons between LiOr and Li_2_CO_3_ (Pacholko & Bekar, 2021). There is very limited data regarding the therapeutic index LiOr in comparison to Li_2_CO_3,_ as current lithium research has only been conducted with equivalent dosages of LiOr and Li_2_CO_3_.

This experiment aimed to help fill the gap in research by systematically comparing LiOr and Li_2_CO_3_ across a range of doses to determine their respective therapeutic indices in terms of memory and locomotive function in a *Drosophila melanogaster* AD model, leading to the research question: how do varying dosages of LiOr and Li_2_CO_3_ compare in improving the locomotion and memory of a *Drosophila melanogaster* model of Alzheimer’s disease?

Given the need to analyze the therapeutic potential of the lithium salts, *Drosophila melanogaster* models provide a potential platform for these needs. *Drosophila melanogaster* models preserve many mammalian processes, including some neurological processes. Not only do *Drosophila melanogaster* share 60% of human genes, specifically having fundamental similarities in the brain, but they also have a rapid generation and a short life span/cycle (Mirzoyan et al., 2019). Their similarity to human brains implies that AD treatments that work on *Drosophila* could work on humans as well, whilst rapid generation and life cycle allow for *Drosophila* to be tested on in a relatively short period, allowing for more trials to be conducted. *Drosophila melanogaster* models are much less complex emotionally than mammalian models, which makes them an ethical model to use for research.

To create a *Drosophila melanogaster* model for AD, the GAL4/UAS system is commonly utilized. This system allows for precise, tissue-specific control of gene expression. In order to use this system, a GAL4 *Drosophila melanogaster* with a specific promoter is maintained and crossed with a *Drosophila melanogaster* stock with a UAS target gene. The GAL4 protein is a transcription factor driven by the promoter. The GAL4 protein will bind to binding sites that are present along the UAS sequence, which means that any gene under the control of the UAS has its transcription activated when GAL4 attaches to the UAS sequence in the tissue location designated by the GAL4, leading to targeted gene expression (Tue et al., 2020).

For this experiment, a UAS stock with the Arctic mutation of Aβ-42 will be used (Bloomington Drosophila Stock Center, n.d.). The Arctic mutation was chosen because it produces more aggressive effects on both locomotion and memory compared to other genes (Prüßing et al., 2013). More aggressive effects of AD will lead to a more distinct comparison of the effects of lithium treatments. An Act5C-GAL4 that expresses GAL4 ubiquitously under the Act5C promoter will be used (Bloomington Drosophila Stock Center, n.d.). (Lee & Koh, 2023). The cross between the Arctic mutation of Aβ-42 and Act5C-GAL4 was chosen due to crosses between Aβ-42 and Act5C-GAL4 leading to locomotive and memory defects in the past (Tan et al., 2023). Therefore, a cross between the Arctic mutation of Aβ-42 and Act5C-GAL4 should be effective for testing the effects of lithium on both locomotion and memory.

It was hypothesized that if a *D. melanogaster* model of AD was subjected to both Li_2_CO_3_ and LiOr treatments across various dosages, LiOr would have a superior therapeutic effect to Li_2_CO_3_ on the locomotion and memory of the AD *D. melanogaster* at lower doses because of its higher bioavailability, allowing for reduced toxicity with lower doses while maintaining neuroprotective benefits. At higher dosages, LiOr is expected to be less effective than Li_2_CO_3_ due to its toxicity at standard Li_2_CO_3_ dosages (Pacholko & Bekar, 2021).

Using varying dosages of LiOr to test for its effect on memory and locomotion may identify advantages of LiOr in mitigating symptoms of AD at lower doses. This experiment will allow us to fill in much-needed gaps in the knowledge of LiOr’s therapeutic index, providing invaluable insight into the viability of LiOr as a potential treatment for AD. If LiOr is found to be a viable alternative to Li_2_CO_3_ to treat locomotive and memory impairments in AD, treatments using LiOr can be developed in order to help improve the quality of life of AD patients.

To examine the locomotive function of *Drosophila*, a negative geotaxis assay will be used. The negative geotaxis assay measures the natural climbing behavior of flies; deficits in this behavior can indicate motor function impairments associated with neurodegeneration. This assay was chosen due to its being very simple, as well as giving a solid understanding of the locomotive capabilities of the flies involved, which will allow for the analysis and comparison of locomotive impacts of lithium salts.

To study memory in *Drosophila*, the aversive phototaxic suppression assay (APS) will be utilized. The APS assay measures the ability of the *Drosophila* to learn and remember to avoid light associated with an aversive stimulus, indicating cognitive function and memory. First, *Drosophila* are trained to associate the aversive smell with light, which goes against their natural phototaxic (attracted to light) behavior. Some time is spent waiting, then their behavior is tested again after some time to see if they remember the association. This assay was selected to measure memory due to the relative ease required to conduct while still providing a good measure of memory and the prevalence of APS in present *Drosophila* research (Torre et al., 2023).

## Materials and Methods

The independent variables were the dosages of either lithium treatment (LiOr and Li_2_CO_3_) in the fly food and the presence of AD in the flies (Act5C-GAL4 *D. melanogaster* vs Alzheimer’s Disease *D. melanogaster* obtained via the crossing scheme). Both lithium drugs were administered at 5 mM and 10 mM to compare multiple studied doses of other lithium drugs (Castillo-Quan, 2016). When converted from mM to mg/mL using the molar masses of LiOr (162.0 g/mol) and Li_2_CO_3_ (73.89 g/mol), these doses correspond to 0.81 mg/mL & 1.62 mg/mL for LiOr and 0.369 mg/mL & 0.739 mg/mL in Li_2_CO_3_. Relevant dosage calculations are shown in Table 2 for LiOr and Table 3 for Li_2_CO_3_.

**Table 1.** Control and Experimental Groups.

| Group | Diet | *D. melanogaster*<br>Model | Purpose for testing | Expected Result on<br>Locomotion and<br>Memory |
| --- | --- | --- | --- | --- |
| Negative<br>Control | Regular<br>Diet | Baseline<br>GAL4 | Provide baseline results<br>for healthy flies | Regular |
| Disease<br>Control | Regular<br>Diet | AD model | Evaluate effectiveness of<br>the AD model | Worse than<br>negative control |
| Li<sub>2</sub>CO<sub>3</sub><br>Toxicity<br>Control 1 | Li<sub>2</sub>CO<sub>3</sub><br>Diet - 5<br>mM | Baseline<br>Gal4 | Evaluate effect of 5 mM<br>Li<sub>2</sub>CO<sub>3</sub> Diet on healthy<br>flies | Same/Worse than<br>negative control |
| Li<sub>2</sub>CO<sub>3</sub><br>Toxicity<br>Control 2 | Li<sub>2</sub>CO<sub>3</sub><br>Diet - 10<br>mM | Baseline<br>Gal4 | Evaluate effect of 10 mM<br>Li<sub>2</sub>CO<sub>3</sub> Diet on healthy<br>flies |  |
| Li<sub>2</sub>CO<sub>3</sub><br>Experimental 1 | Li<sub>2</sub>CO<sub>3</sub><br>Diet - 5<br>mM | AD Model | Evaluate 5 mM Li<sub>2</sub>CO<sub>3</sub><br>Diet's effect in AD model | Better than disease<br>control |
| Li<sub>2</sub>CO<sub>3</sub><br>Experimental 2 | Li<sub>2</sub>CO<sub>3</sub><br>Diet - 10 | AD Model | Evaluate 10 mM Li<sub>2</sub>CO<sub>3</sub><br>Diet's effect in AD model | Better than the 10<br>mM LiOr |

|  | mM |  |  | experimental group |
| --- | --- | --- | --- | --- |
| LiOr Toxicity<br>Control 1 | LiOr Diet<br>- 5 mM | Baseline<br>Gal4 | Evaluate effect of 5 mM<br>LiOr Diet on healthy flies | Same/Worse than<br>negative control |
| LiOr Toxicity<br>Control 2 | LiOr Diet<br>- 10 mM | Baseline<br>Gal4 | Evaluate effect of 10 mM<br>LiOr Diet on healthy flies |  |
| LiOr<br>Experimental 1 | LiOr Diet<br>- 5 mM | AD Model | Evaluate 5 mM LiOr<br>Diet's effect in AD model | Better than disease<br>control |
| LiOr<br>Experimental 2 | LiOr Diet<br>- 10 mM | AD Model | Evaluate 10 mM LiOr<br>Diet's effect in AD model | Same/Worse than<br>the 10 mM Li<sub>2</sub>CO<sub>3</sub><br>experimental group |

*Note.* This table presents the structure and function of the groups. Groups filled in RGB (220, 203, 226) are AD Model groups.

**Figure 1.**
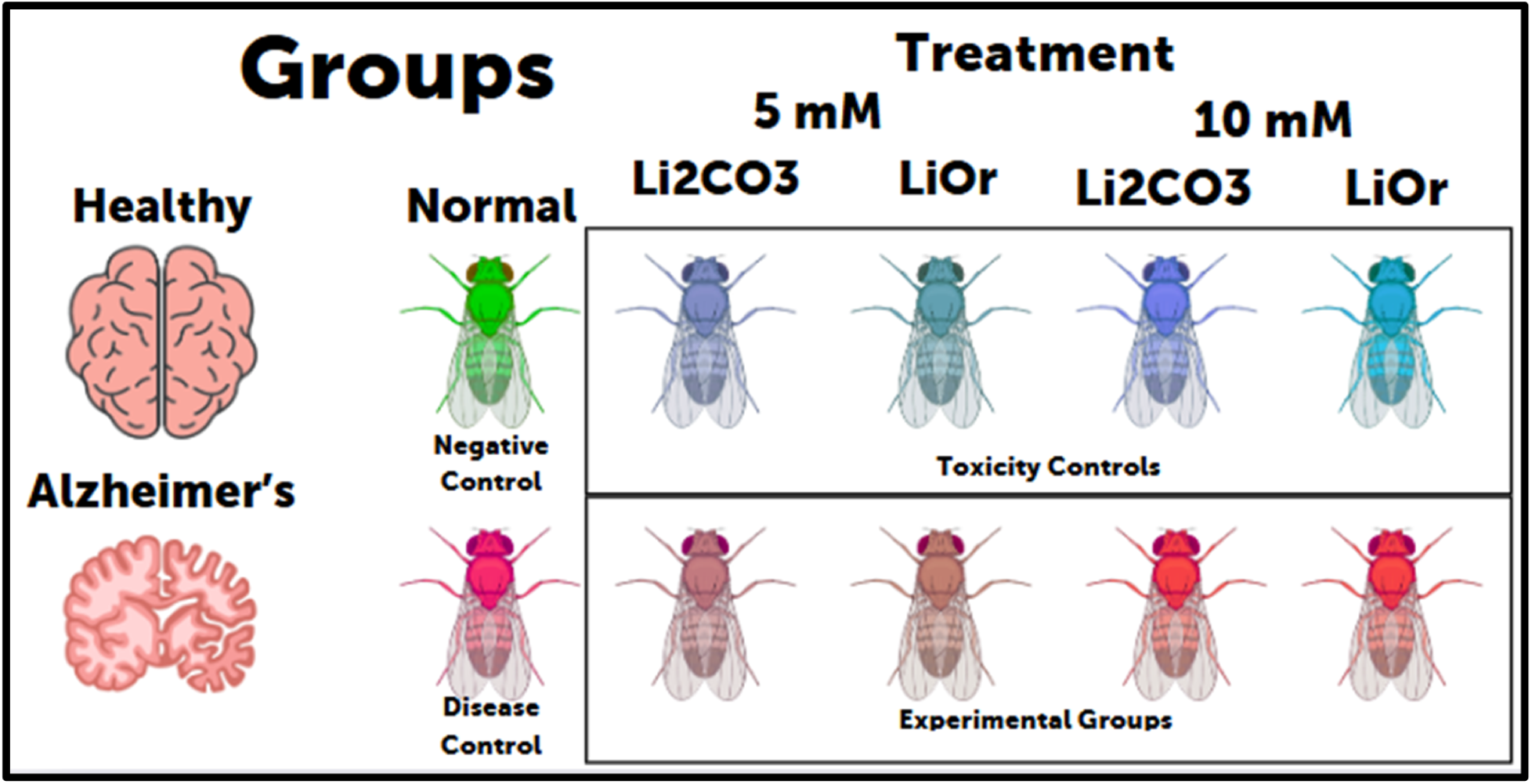
Groups. *Note.* This figure presents a diagram of the groups compared.

The dependent variables were locomotive function as measured using the negative geotaxis assay by the percentage of flies that successfully climb to a specified height in 10 seconds, and memory as measured using the Aversive Phototaxic Suppression (APS) assay by the percentage of flies that go toward the dark vial.

Constants for this experiment included the age of flies (28 days), the time of data collection (1:50 PM - 3:30 PM EST), the sex of flies (male), the lab equipment used, and the location (Dr. Eliason’s Lab - Academies of Loudoun Room 2419).

This project included six control groups (One negative control, one disease control, two Li_2_CO_3_ toxicity controls and two LiOr toxicity controls for 5 and 10 mM) and four experimental groups (two LiOr treated AD and two Li_2_CO_3_ treated AD for 5 and 10 mM). The negative control group will use Act5c-GAL4 *D. melanogaster* with no AD present and will not be treated with any lithium salts. This group provides insight into what the expected results of the assays are, allowing for the comparison of experimental groups with healthy flies. The negative control should perform well in both the climbing and memory assay due to there being no AD present. It should be noted that parent flies, flies without AD, will be used rather than wildtype, as transgenic flies may perform differently than wildtype. The disease control group will demonstrate the effects of the AD model being used. The disease control should perform worse than the negative control, because of the presence of AD. The presence of the disease control allows for the determination of whether or not the AD model works, or if either lithium salt makes any significant improvement over the base AD *Drosophila*.

The other control groups are the toxicity control groups, which consist of *D. melanogaster* without AD present that are treated with Li_2_CO_3_ or LiOr. These control groups provide insight on the effects of Li_2_CO_3_ and LiOr on healthy flies, which is important specifically for this project due to the negative side effects associated with lithium salt treatments. These control groups are expected to perform slightly worse than the negative control group due to the aforementioned negative side effects of lithium salt treatments.

### Materials

[AOS] indicates that the material will be sourced from the Academy of Science, rather than purchased online.

#### Flies

- *D. melanogaster* Act5C-GAL4 [Bloomington *Drosophila* Stock Center - Stock #4414]
- *D. melanogaster* UAS-APP.Abeta42.E693G [Bloomington *Drosophila* Stock Center - Stock #33774]
- Fly Storage [AOS]
  ○ Sharpie
  ○ Tape
  ○ Clear plastic vials
  ○ Foam plugs (flugs)
- Fly Food
  ○ Yellow Corn Meal [AOS]
  ○ Distilled water [AOS]
  ○ Light corn syrup [AOS]
  ○ Soy flour [AOS]
  ○ Yeast [AOS]
  ○ Agar [AOS]
  ○ Li_2_CO_3_ [AOS]
    ▪ https://www.fishersci.com/store/msds?partNumber=L119500+productDescription=LITHIUM+CARBONATE+CR+ACS+500G+vendorId=VN00033897+countryCode=US+language=en
  ○ Propionic acid - (10%, AOS)
    ▪ https://www.fishersci.com/store/msds?partNumber=A258500+countryCode=US+language=en
  ○ LiOr (betterLife.com, 2019)
    ▪ https://www.chemicalbook.com/msds/Lithium-orotate.pdf
  ○ Cheesecloth [AOS]

### Lab Equipment [AOS]

- Paper towels
- Thermometer
- Pipette
- Graduated cylinders (100mL, 500mL)
- Beaker (500 mL, 1L)
- Weigh boats
- Scales
- Freezer
- Scoopulas
- Microwave (Panasonic 1250W Genius Sensor)
- Funnel
- Glass stirring rod

### Sorting Equipment [AOS]

- Weigh paper
- Magnifying glass
- Feather
- Cold sorter (AGP-301CPV)
- CO_2_ plate
- Arc3 gasses CO_2_ tank
- Genesee scientific Flystuff Blowgun
- Labomed Microscope Luxeo 6z
- Ice
- Ice Bucket

### Climbing Assay Equipment [AOS]

- Cotton balls
- Graduated cylinder (100mL)
- Marker
- Ruler
- Sharpie
- Tape
- Timer
- Video recording device

### APS Assay [AOS]

- ≥95%, FG Quinine Hydrochloride Dihydrate
  ○ https://www.fishersci.com/store/msds?partNumber=AAH33474+productDescription=keyword+vendorId=VN00024248+countryCode=US+language=en
- Clear plastic vials
- Index card
- 0.45 mm Filter paper
- Fume hood
- LED fairy lights
- Opaque black tape
- Timer

### Safety Equipment [AOS]

- Oven Mitts
- Goggles
- Lab Coats
- Closed-toe shoes

### Fly food preparation

Before experimentation, food for the treated and untreated groups was created. To prepare food, a weighboat and scoopula were used to measure out 6.75 g of yeast, 3.90 g of flour, 28.50 g of yellow cornmeal, and 2.25 g of agar and combined with 390 mL of distilled water (H_2_O) and 30 mL of light corn syrup into a 1L beaker using graduated cylinders and the scale. A glass stirring rod was used to mix the contents of the beaker until all materials were dissolved. The mixture was then placed into the microwave, where it was heated in 1-minute intervals until the mixture bubbled and rose. Following each minute, the beaker was removed from the microwave and stirred with the stirring rod. This process was repeated 3 times per batch of food made. Cheesecloth was put over the mixture and the mixture was cooled to 70°F. Once at 70°F, a micropipette was used to add 1.88 mL of propionic acid to the mixture and then mixed in using a stirring rod. Once stirred, about 10 mL of the food was placed into each empty vial, each vial was labelled, and cheesecloth was placed on the vials. Once all vials were filled, a flug was then placed on each vial and the vial tray was put into the fridge until use.

Food for treated groups was produced using the same procedure mentioned above, with amounts of LiOr and Li_2_CO_3_ added into the beaker and dissolved. For LiOr, 375 mg was added for 5 mM and 751 mg was added for 10 mM as calculated in Table 2. For Li_2_CO_3_, 171 mg was added for 5 mM and 342 mg was added for 10 mM as calculated in Table 3.

**Table 2.** LiOr Diet Preparation Calculations.

| Concentration (mM) | 5 | 10 |
| --- | --- | --- |
| Conversion to mg/mL | 5 mmol/L LiOr x 162.0<br>mg/mmol x 1L/1000mL | 10 mmol/L LiOr x 162.0<br>mg/mmol x 1L/1000mL |
| Concentration (mg/mL) | 0.810 | 1.62 |
| LiOr Mass Calculation | 463.28 mL x 0.810 mg/mL =<br>375 mg | 463.28 mL x 1.62 mg/mL =<br>751 mg |

*Note.* This table presents the calculations for the amount of LiOr to add to the food mixture.

**Table 3.** Li_2_CO_3_ Diet Preparation Calculations.

| Concentration (mM) | 5 | 10 |
| --- | --- | --- |
| Conversion to mg/mL | 5 mmol/L Li<sub>2</sub>CO<sub>3</sub> x 73.89<br>mg/mmol x 1L/1000mL | 10 mmol/L Li<sub>2</sub>CO<sub>3</sub> x 73.89<br>mg/mmol x 1L/1000mL |
| Concentration (mg/mL) | 0.369 | 0.739 |
| Li<sub>2</sub>CO<sub>3</sub> Mass Calculation | 463.28 mL x 0.369 mg/mL =<br>171 mg | 463.28 mL x 0.739 mg/mL =<br>342 mg |

*Note.* This table presents the calculations for the amount of Li<sub>2</sub>CO<sub>3</sub> to add to the food mixture.

### Sorting flies

Two different sorting methods were used, carbon dioxide (CO_2_ sorting) and cold sorting. CO_2_ sorting was used to sort flies that would be used for crosses, whilst cold sorting was used to sort flies that would be used for experimentation due to the damage that CO_2_ can cause to the brain of flies.

To sort using CO_2_, firstly the CO_2_ canister and CO_2_ sorter lights had to be turned on so that CO_2_ was flowing. Once both were turned on, the fly vials were flipped upside down and tapped so that the flies were on the flug. After the flies were on the flug, the CO_2_ gun was inserted into the side of the flug and CO_2_ flowed into the vial, causing the flies to fall asleep. Consequently, once the flug was removed, the flies would fall out of the vial and onto the CO_2_ sorter, which worked as a pad that would keep the flies asleep. On the pad, the flies remained asleep, allowing the usage of a microscope and feather to sort through the flies and group them based on characteristics (sex, wing shape, virginity, etc.). The flies were placed into new vials facing sideways using the feather based on the traits sorted, and the CO_2_ was turned off.

To sort using cold sorters, the following procedure was used. First, the cold sorter was turned on and set to 0°C. While it cooled to that temperature, flies that needed to be sorted were tapped into empty in an ice bath for 3-4 minutes, or until the flies were asleep (checked every 30 seconds). Once asleep, the flies were placed on the cold sorter plate to ensure they would not wake up. A magnifying glass was used to determine the traits of the flies and a feather was used to move the flies into their respective groups on the cold sorting plate. After grouping the flies based on traits, the flies were tapped back into vials facing sideways using the feather.

To determine the sex of the flies, the visual appearance of the fly was analyzed. Female flies are lighter, larger, have more stripes on the abdomen, a pointed tip, no sex combs or claspers, whilst males flies have claspers, sex combs, and a round, dark tip, while being much smaller than the female. The microscope or magnifying glass was used in order to figure out which traits a given fly matches most closely with, allowing for the identification of the sex of the fly. To identify virgin females, a closer examination of the flies took place. Virgin flies have larger, pale bodies, with folded wings or the presence of a meconium patch, guaranteeing the virginity of the fly, whilst the presence of sex combs was used to determine sex.

### Fly stocks

Two stocks were obtained from Bloomington *Drosophila* Stock Center at Indiana University: Act5C-GAL4 (stock #4414) & UAS-APPAbeta42.E693G (stock #33774). When obtained, the stocks were put into vials with fly food.

To tap or transfer files to a different vial, a vial with flies in it and a vial with fresh food were gathered. With one vial in each hand, the flug was removed from the vial with fresh food. The new vial was held upside down and the vial with flies was tapped ∼2-3 times until the flies were at the bottom. The flug was removed from the vial with flies and the fresh vial was placed on it, ensuring there was no space for the flies to escape. The two vials were then flipped upside down while maintaining contact, and then tapped onto the table to drop the flies to the bottom of the fresh vial. New flugs were then placed on the fresh and old vial while keeping the old vial upside down. The new vial was then labelled.

### Crossing stocks

In order to cross stocks to obtain progeny modeling AD, 5-6 virgin females of UAS and 4 males of GAL4 were gathered as indicated in the crossing scheme in Tables 4 & 5 and Figure 2. Because new flies hatch in 14 days and can reproduce, parent flies were tapped out before hatching. *Drosophila* with curly wings (CyO marker) were removed to ensure only heterozygous AD model flies were tested as modeled in Table 4, Table 5, and Figure 2.

**Table 4.** AD D. melanogaster model Punnett Square for Chromosome 3.

|  |  |  |
| --- | --- | --- |
|  | + | + |
| UAS-APP.Abeta42.E693G | UAS-APP.Abeta42.E693G/+ | UAS-APP.Abeta42.E693G/+ |
| UAS-APP.Abeta42.E693G | UAS-APP.Abeta42.E693G/+ | UAS-APP.Abeta42.E693G/+ |

*Note.* This table presents the Punnett Square for Chromosome 3 for the crossing scheme used.

**Table 5.**
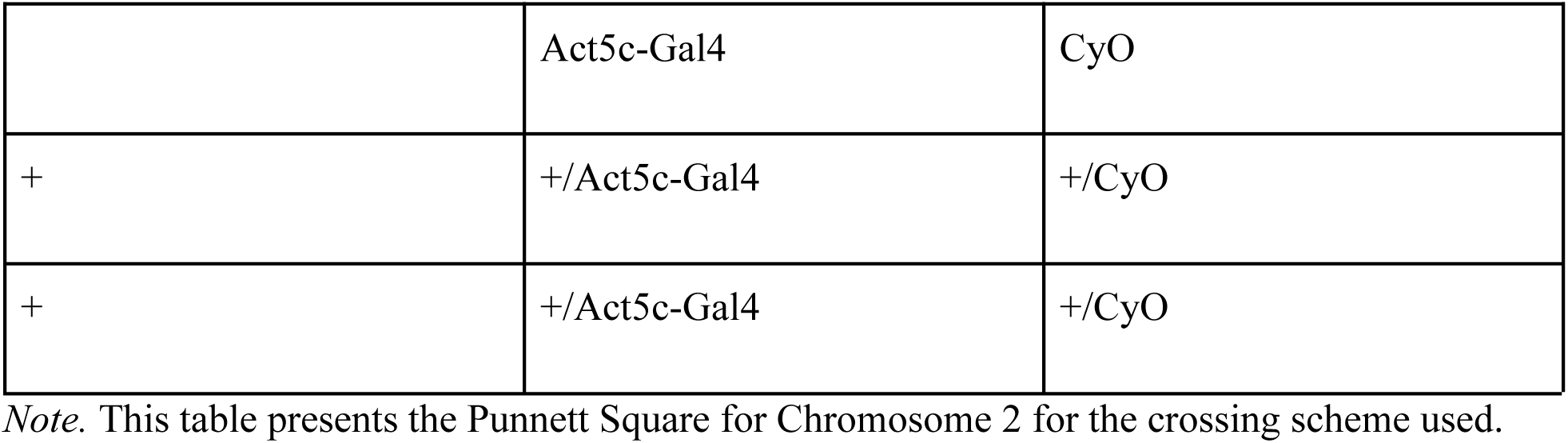
AD D. melanogaster model Punnett Square for Chromosome 2.

**Figure 2.**
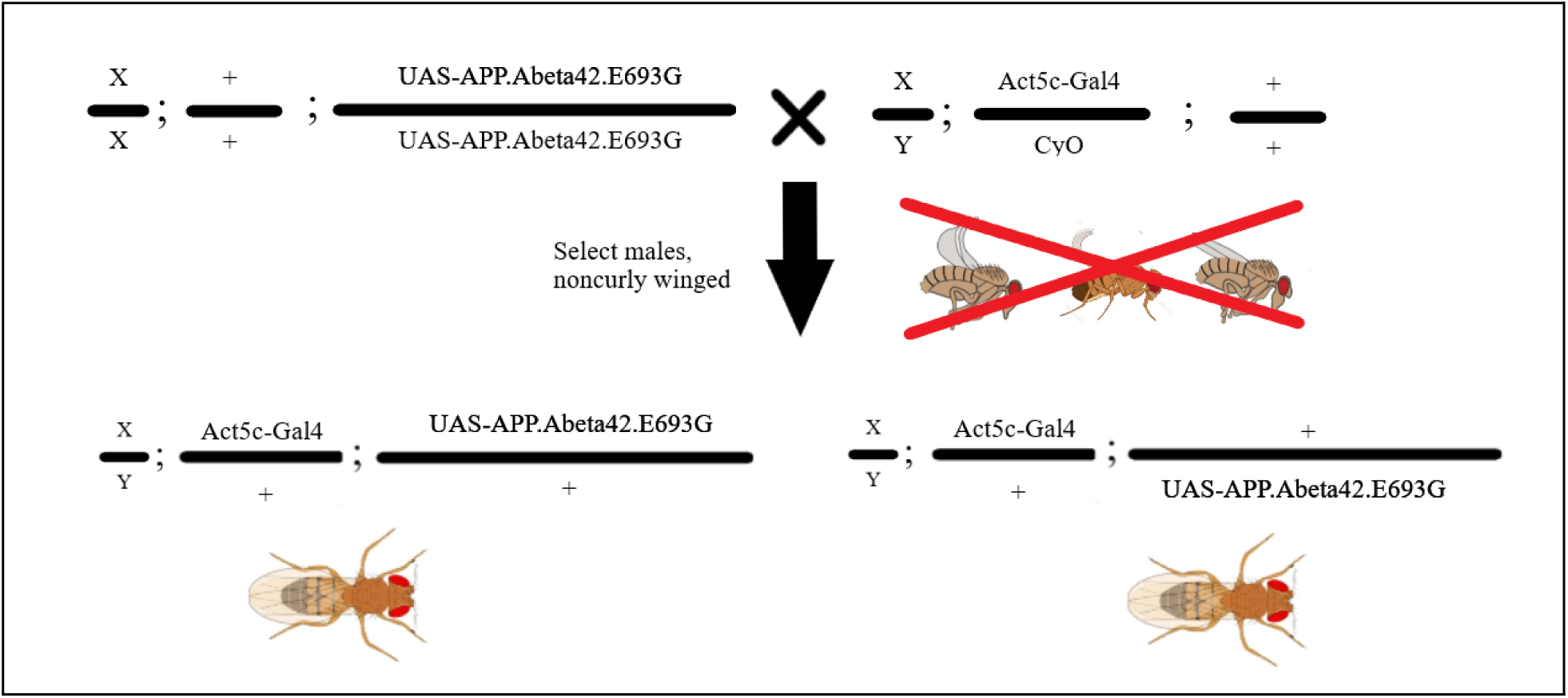
AD D. melanogaster model crossing scheme. *Note.* This figure presents a design of the crossing scheme performed.

### Negative geotaxis (Climbing) assay

The climbing assay was used to determine the locomotive capability of the flies tested. To begin, male flies were cold sorted into groups of 10 flies each. For the apparatus, a rule was used to mark 8 cm above the bottom of a graduated cylinder, which was marked using a piece of tape. Flies were tapped into the graduated cylinder, which was sealed using a cotton ball. A video recording device (e.g., a phone) was used to record each trial, ensuring the tape and flies were visible, as well as a stopwatch as well. The recording was started along with the stopwatch, and the flies were tapped to the bottom of the graduated cylinder. The flies were given 10 seconds to climb along the graduated cylinder, then disposed of. The video was then analyzed to determine the total number of flies that crossed the tape (passed), using the time seen on the stop watch when the flies are tapped as the start time, and 10 seconds after as the end time. It should be noted that each fly could only count once, once a fly crossed it counted no matter where it finished the assay, and flies had to fully cross the 8cm mark. The pass rate for each trial was then calculated using the formula: Pass rate (%) = (Number of passes/total number of flies) * 100%.

**Figure 3.**
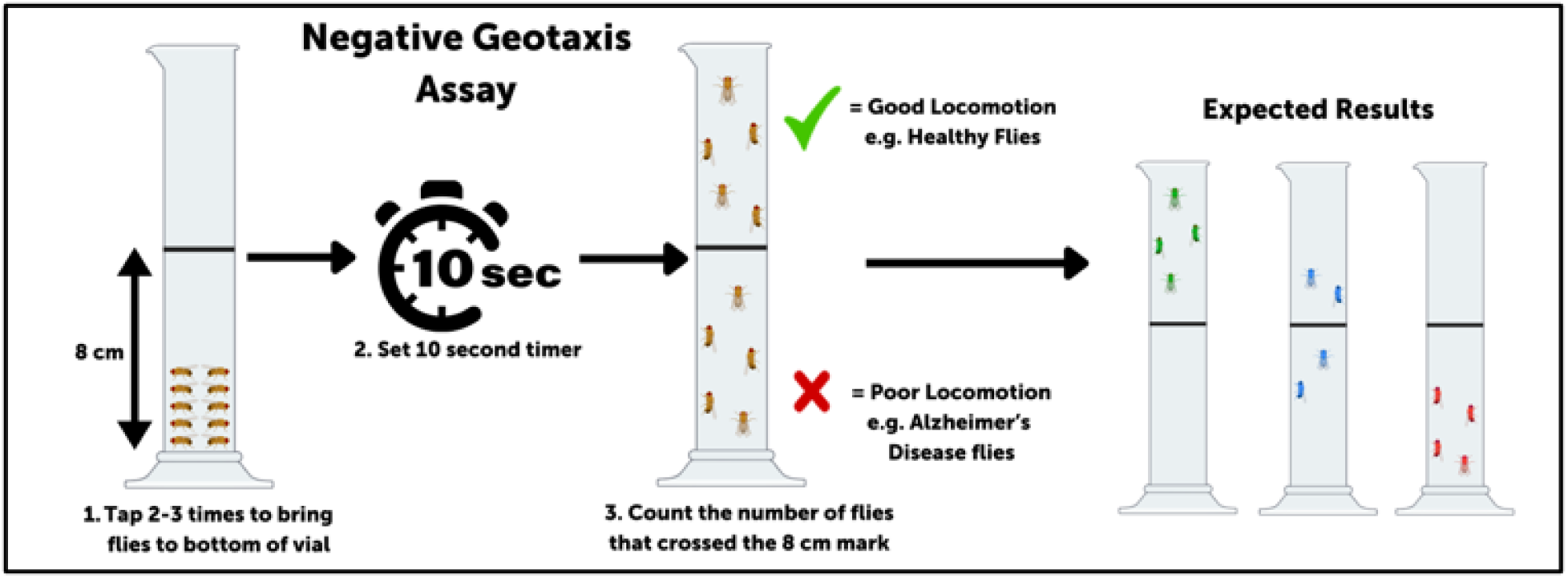
Climbing/Negative Geotaxis assay diagram. *Note.* This figure presents a diagram of the negative geotaxis (climbing) assay used.

### Aversive phototaxic suppression (APS) assay

The APS assay was used to determine the memory capability of the flies tested. First, solid quinine hydrochloride was diluted under a fume hood to create 200μL of 0.1 M quinine hydrochloride by the Chemical and Laboratory Safety Specialist (dissolve 7.9382mg of solid quinine hydrochloride in 200μL of water; 0.1mmol/mL x 0.2 mL * 396.91 mg/mmol = 7.9382mg). Then, male flies were cold sorted into groups of 10 each. Two vials with a flug in them were made, one of which was covered in opaque black tape surrounding it, and the other had fairy lights surrounding it. In the vial with lights, 200μL of 0.1 M quinine hydrochloride was put into the flug using a micropipette. To train the flies, the group of 10 flies was tapped into the dark vial, where they were allowed to acclimate for 30 seconds. After, they were tapped into the light vial for another 30 seconds. The flies were then tapped back and forth for a total of 10 cycles. An hour after training, flies were tapped into the middle of two odorless vials (one dark & one light), and given 30 seconds to go towards either side. If the flies ended up in the dark vial, it was considered a pass, while if a fly ended up in the light vial it was considered a fail. The pass rate of the group was then calculated using the same formula as the climbing assay.

**Figure 4.**
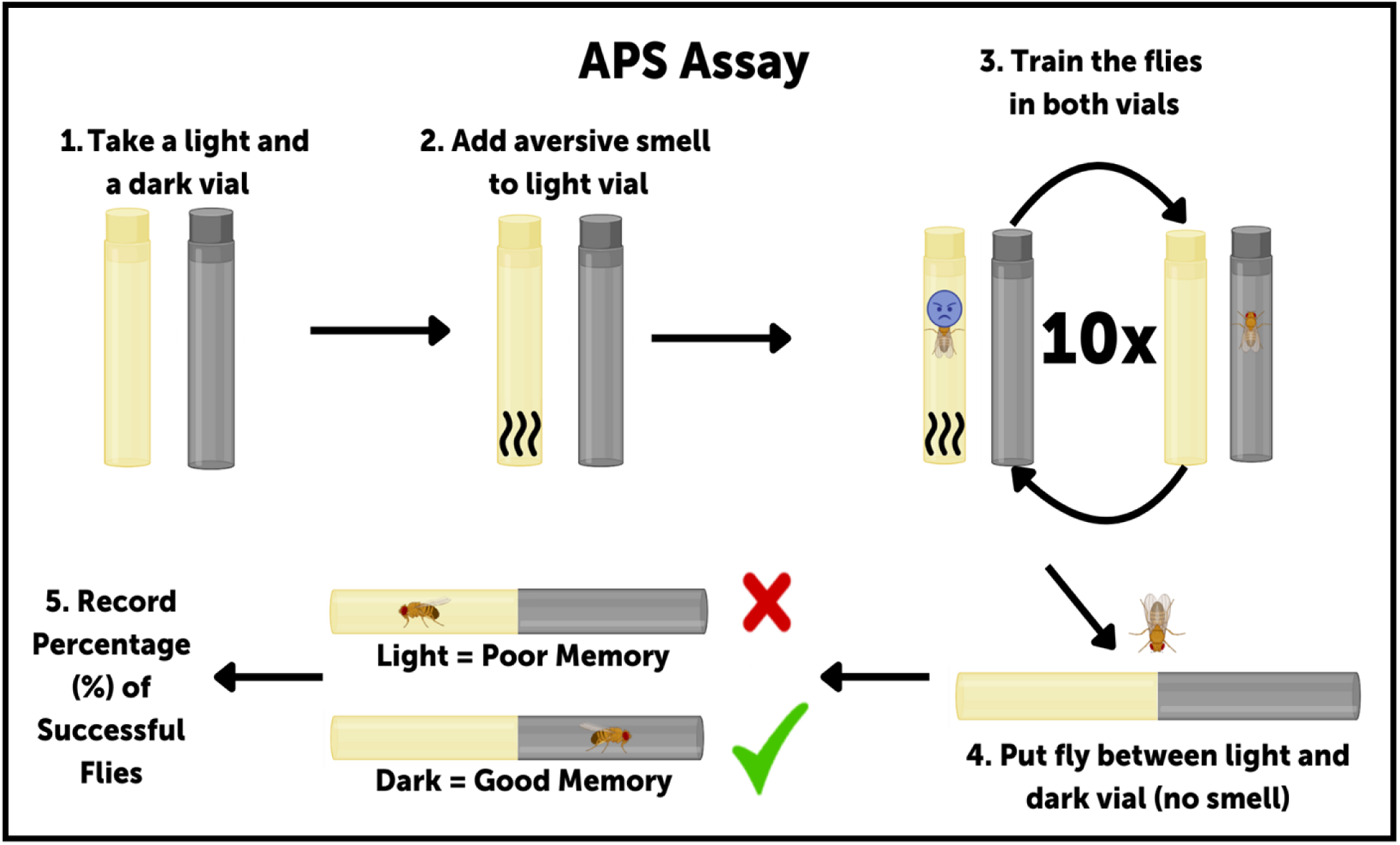
Aversive phototaxic suppression (APS) assay diagram. *Note.* This figure presents a diagram of the aversive phototaxic suppression (APS) assay used.

### Statistical analysis

Post data collection, statistical analysis was conducted to determine if there was any significant difference between groups. Because a normal distribution could not be confirmed due to a sample size less than 30 samples, nonparametric statistical tests were used. The Mann-Whitney U test was used when comparing APS and negative geotaxis assay pass rates between healthy and AD flies & Li_2_CO_3_ and LiOr at 5 mM and 10 mM. The Kruskal-Wallis test was used when comparing APS and negative geotaxis assay pass rates between healthy flies, 5 mM lithium drug healthy flies, & 10 mM lithium drug healthy flies for both LiOr and Li_2_CO_3_ within both healthy flies and AD flies.

### Safety

Throughout the experiment, certain general safety procedures took place to ensure no injuries or accidents occurred from experimentation. To begin, the experiment occurred in a clear, safe environment where closed-toed shoes were worn. Personal Protective Equipment (PPE) of safety goggles and a lab coat was worn to protect the eyes from glassware (hazard if it breaks) and skin from chemical exposure when either glassware or chemicals that required it were utilized. Hands were also washed with soap and water thoroughly before and after the experiment. Heat-resistant mitts were worn when boiling fly food to protect skin from the heat.

Specific chemicals also had safety protocols associated with them as well. For propionic acid these protocols included keeping the acid away from sparks, heat, open flames, and hot surfaces as propionic acid is flammable, storage in a well-ventilated area, and the disposal of pipette tips or anything else in contact with propionic acid in special waste disposal units or bags.

For Li_2_CO_3_, these protocols included storage in a dry, cool, well-ventilated area. The avoidance of inhalation of mist gas or vapors and dust collected nearby was followed, as lithium could remain on the dust particles. Disposal of anything that came in contact with the lithium in a special waste disposal unit or bag and discharge in the environment were followed. The procedures for LiOr were the same as the protocols for Li_2_CO_3_.

For quinine hydrochloride, these protocols included the avoidance of inhalation or ingestion, no dust being created nearby, the avoidance of sparks, heat, open flames, and hot surfaces due to quinine being combustible, and the disposal of anything that came in contact with the quinine in waste disposal units or bags.

These protocols, along with the general safety measures listed before, were followed in order to avoid dangerous events. If an event had happened, the lab supervisor would have been contacted along with any emergency personnel, the MSDS protocols would have been followed given the circumstances, and first aid would be administered accordingly to the accident.

### Data

**Table 5.** Climbing Assay Pass Rate Data.

| Successful flies in Climbing Assay (%) |  |  |  |  |  |  |  |  |  |  |
| --- | --- | --- | --- | --- | --- | --- | --- | --- | --- | --- |
| Groups | Normal<br>Gal4 | 5mM<br>LiOr<br>Gal4 | 5mM<br>Li<sub>2</sub>CO<sub>3</sub><br>Gal4 | 10mM<br>LiOr<br>Gal4 | 10mM<br>Li<sub>2</sub>CO<sub>3</sub><br>Gal4 | Normal<br>AD | 5mM<br>LiOr<br>AD | 5mM<br>Li<sub>2</sub>CO<sub>3</sub><br>AD | 10mM<br>LiOr<br>AD | 10mM<br>Li<sub>2</sub>CO<sub>3</sub><br>AD |
| Trial 1 | 71.43 | 44.44 | 70.00 | 63.64 | 66.67 | 33.33 | 40.00 | 44.44 | 44.44 | 70.00 |
| Trial 2 | 63.64 | 80.00 | 81.82 | 72.73 | 80.00 | 27.27 | 45.45 | 33.33 | 42.86 | N/A |
| Trial 3 | 80.00 | N/A | 66.67 | N/A | N/A | N/A | N/A | N/A | N/A | N/A |
| Trial 4 | 81.82 | N/A | 55.56 | N/A | N/A | N/A | N/A | N/A | N/A | N/A |
| Trial 5 | N/A | N/A | 71.43 | N/A | N/A | N/A | N/A | N/A | N/A | N/A |
| **Average** | **74.22** | **62.22** | **69.09** | **68.18** | **73.33** | **30.30** | **42.73** | **38.89** | **43.65** | **70.00** |

*Note.* This table presents the raw data collected from the climbing assays conducted in this project by pass rate (%). “N/A” indicates that the trial was not conducted due to fly mortality or experimental constraints.

**Table 6.** APS Assay Pass Rate Data.

| Successful flies in APS Assay (%) |  |  |  |  |  |  |  |  |  |  |
| --- | --- | --- | --- | --- | --- | --- | --- | --- | --- | --- |
| Groups | Normal Gal4 | 5mM LiOr Gal4 | 5mM Li<sub>2</sub>CO<sub>3</sub> Gal4 | 10mM LiOr Gal4 | 10mM Li<sub>2</sub>CO<sub>3</sub> Gal4 | Normal AD | 5mM LiOr AD | 5mM Li<sub>2</sub>CO<sub>3</sub> AD | 10mM LiOr AD | 10mM Li<sub>2</sub>CO<sub>3</sub> AD |
| Trial 1 | 80.00 | 72.73 | 66.67 | 81.82 | 71.43 | 12.50 | 44.44 | 40.00 | 45.45 | 55.56 |
| Trial 2 | 90.91 | 81.82 | 66.67 | 72.73 | 80.00 | 71.43 | 44.44 | 44.44 | 50.00 | N/A |
| Trial 3 | 87.50 | N/A | 63.64 | N/A | N/A | 11.11 | N/A | N/A | N/A | N/A |
| Trial 4 | 83.33 | N/A | N/A | N/A | N/A | 28.57 | N/A | N/A | N/A | N/A |
| Trial 5 | 88.89 | N/A | N/A | N/A | N/A | 22.22 | N/A | N/A | N/A | N/A |
| **Average** | **86.13** | **77.27** | **65.66** | **77.27** | **75.71** | **29.17** | **44.44** | **42.22** | **47.73** | **55.56** |

*Note.* This table presents the raw data collected from the APS assays by pass rate (%). N/A means no data was collected as that trial was not conducted.

## Results

**Figure 5.**
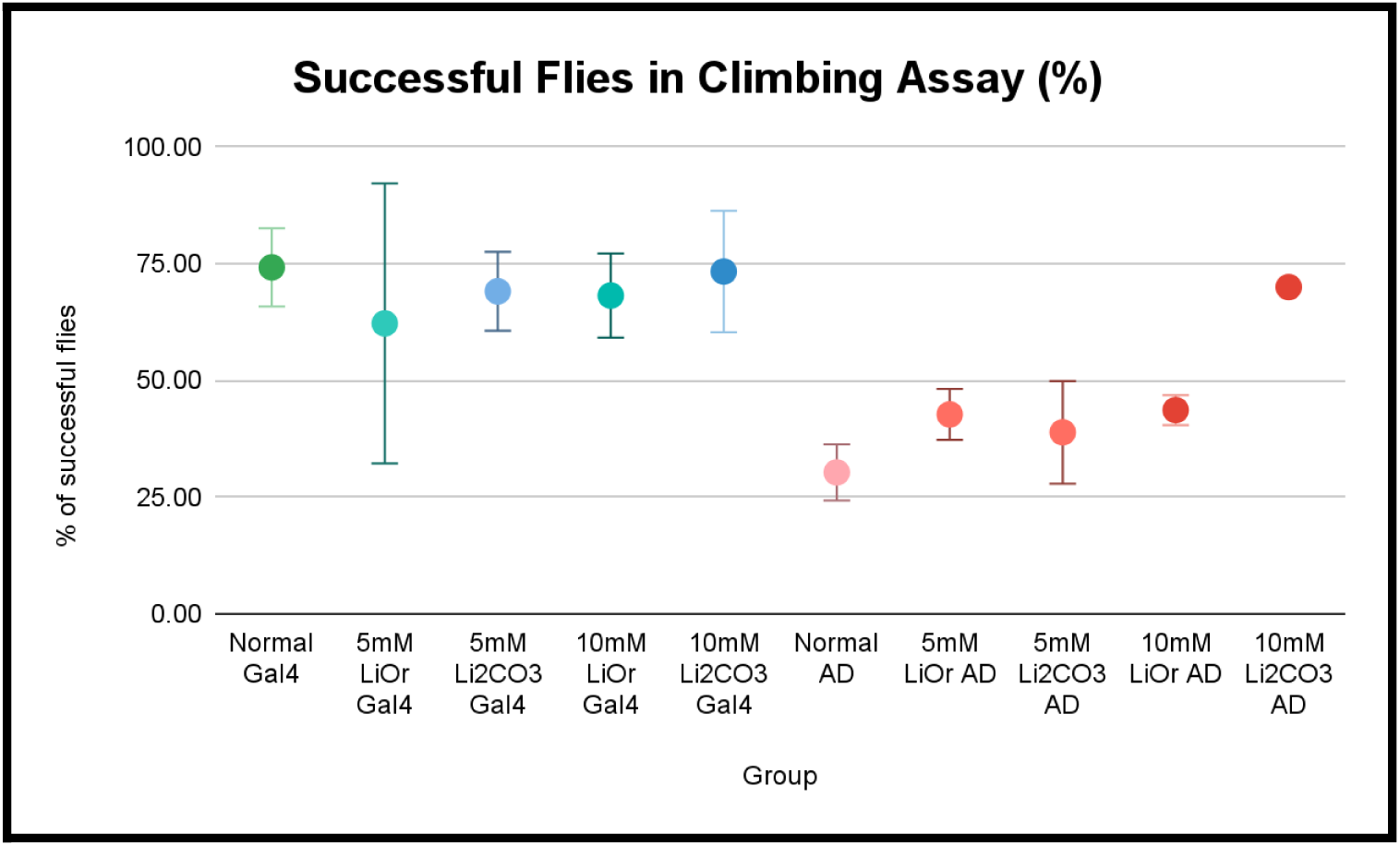
Climbing Assay Pass Rate Graph. *Note.* This graph presents the data collected from the climbing assays by pass rate %. Error bars represent ± 2 SE.

**Figure 6.**
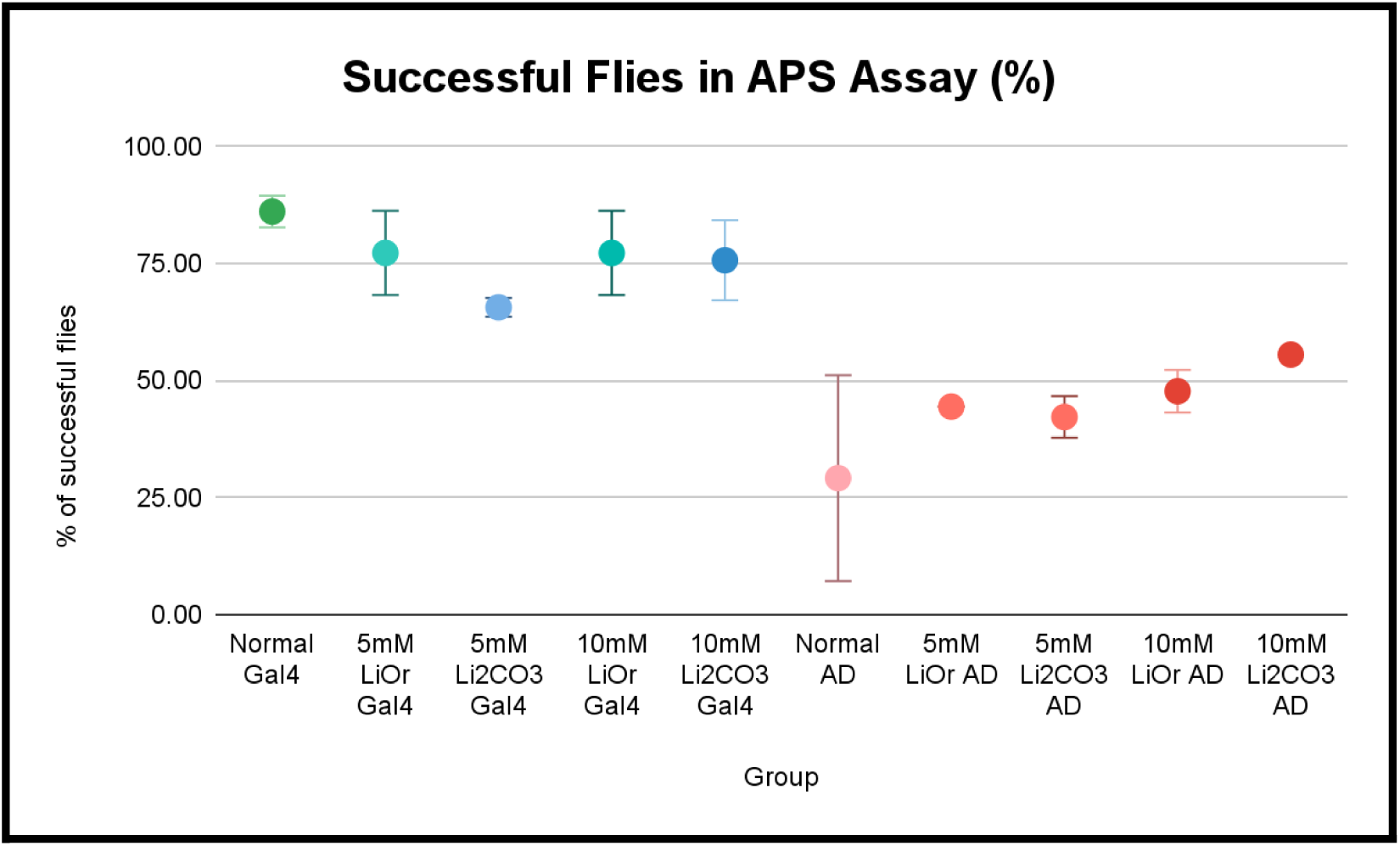
APS Assay Pass Rate Graph. *Note.* This graph presents the data collected from the climbing assays conducted by pass rate %. Error bars represent ± 2 SE. The standard deviation and standard error of each group were also calculated.

**Table 7.** Standard deviation results from each group in each assay.

| Standard Deviation of Pass Rate in Assays (%) |  |  |  |  |  |  |  |  |  |  |
| --- | --- | --- | --- | --- | --- | --- | --- | --- | --- | --- |
| Groups | Normal Gal4 | 5mM LiOr Gal4 | 5mM Li<sub>2</sub>CO<sub>3</sub> Gal4 | 10mM LiOr Gal4 | 10mM Li<sub>2</sub>CO<sub>3</sub> Gal4 | Normal AD | 5mM LiOr AD | 5mM Li<sub>2</sub>CO<sub>3</sub> AD | 10mM LiOr AD | 10mM Li<sub>2</sub>CO<sub>3</sub> AD |
| Climbing Assay | 8.39 | 25.14 | 9.45 | 6.43 | 9.43 | 4.29 | 3.86 | 7.86 | 1.12 | N/A |
| APS Assay | 4.41 | 6.43 | 1.75 | 6.43 | 6.06 | 24.69 | 0.00 | 3.14 | 3.21 | N/A |

*Note.* This graph presents the standard deviation calculated for each group in each assay using the stdev function on excel. Values of N/A represent that there was not enough data in that group to calculate a standard error.

**Table 8.** Standard error results from each group in each assay.

| Standard Error of Pass Rate in Assays (%) |  |  |  |  |  |  |  |  |  |  |
| --- | --- | --- | --- | --- | --- | --- | --- | --- | --- | --- |
| Groups | Normal Gal4 | 5mM LiOr Gal4 | 5mM Li<sub>2</sub>CO<sub>3</sub> Gal4 | 10mM LiOr Gal4 | 10mM Li<sub>2</sub>CO<sub>3</sub> Gal4 | Normal AD | 5mM LiOr AD | 5mM Li<sub>2</sub>CO<sub>3</sub> AD | 10mM LiOr AD | 10mM Li<sub>2</sub>CO<sub>3</sub> AD |
| Climbing Assay | 4.19 | 17.78 | 4.22 | 4.55 | 6.67 | 3.03 | 2.73 | 5.55 | 0.80 | N/A |
| APS Assay | 1.97 | 4.55 | 1.01 | 4.55 | 4.29 | 11.04 | 0.00 | 2.22 | 2.28 | N/A |

*Note.* This graph presents the standard error calculated for each group in each assay using the deviations seen in Table 7. Values of N/A represent that there was not enough data in that group to calculate a standard error.

**Table 9.** Nonparametric Statistical Testing Results.

| Nonparametric statistical testing results |  |  |
| --- | --- | --- |
| Comparison | APS Assay *p* value | Negative Geotaxis Assay *p* value |
| Healthy vs. AD | 0.0121 | 0.405 |
| 0 mM, 5 mM, and 10 mM LiOr | 1.0000 | 1.0000 |
| 0 mM, 5 mM, and 10 mM Li<sub>2</sub>CO<sub>3</sub> | 0.0434 | 0.6121 |
| 5 mM Li<sub>2</sub>CO<sub>3</sub> vs. LiOr | 0.1421 | 0.8631 |
| 10 mM Li<sub>2</sub>CO<sub>3</sub> vs. LiOr | 0.6742 | 0.6742 |
| 0 mM, 5 mM LiOr, and 10 mM LiOr in AD | 0.2615 | 0.1824 |
| 0 mM, 5 mM, and 10 mM Li<sub>2</sub>CO<sub>3</sub> in AD | 0.2185 | 1.0000 |
| 5 mM Li<sub>2</sub>CO<sub>3</sub> vs. LiOr in AD | 0.6251 | 0.6666 |
| 10 mM Li<sub>2</sub>CO<sub>3</sub> vs. LiOr in AD | 0.6685 | 0.6685 |

*Note.* This table presents the *p* values from the nonparametric tests used as described earlier.

## Discussion/Analysis

A difference was found between the memory of control groups of healthy flies and healthy flies treated with Li_2_CO_3_ for the APS assay (*p* = 0.0121), but this difference was not present for the negative geotaxis assay (*p* = 0.405). The difference in the APS assay supports previous research that lithium can improve cognitive function at lower doses in animal models and humans (Forlenza et al., 2014; Shin et al., 2023). The control for Li_2_CO_3_’s effectiveness in healthy flies was similarly effective in the APS assay (*p* = 0.0434), but not effective for the negative geotaxis assay (*p* = 0.6121). Locomotion yielding insignificant results while memory yields significant results in both controls highlights a likely inaccuracy of data due to low sample size. Because comparisons for experimental groups failed to yield significant results with an alpha level of 0.05, it was concluded that Li_2_CO_3_ or LiOr did not significantly affect the locomotion of healthy flies or the locomotion or memory of AD flies. The comparisons between other control groups highlights the lack of data within this project or procedural errors.

While past studies (Kling et al., 1978; Pacholko & Bekar, 2021) suggested that LiOr’s higher bioavailability could lead to greater neuroprotective effects at lower dosages, the findings did not support a significant advantage of LiOr over Li_2_CO_3_ at the tested dosages, which may have resulted from differences in experimental design, dosage range, or the specific AD model used. Because other assays such as the lifespan and toxicity assays were not used, the optimal dosage of the lithium drugs was unclear with full certainty. Rather than using less mainstream, aggressive models like the ‘Arctic’ AD model used, a more researched GAL4-UAS model should be prioritized for more consistent controls to highlight the effectiveness of treatments. Using an Alzheimer’s Disease model without a crossing scheme necessary would significantly decrease the time needed for trials and could increase the sample size.

A notable limitation was the low sample size and resulting high variability, as reflected in the error values in Tables 7 and 8. The high standard deviation and standard error are likely due to the inconsistencies in the data that resulted from the variance associated with small samples. Higher standard deviations also leads to less significant differences as the variance is higher, providing a possible explanation for the lack of statistical significance found.

A major hindrance was the weather conditions that hindered the research progress in January, along with a fire at the school causing closure. These closures meant that for a portion of the year, students were not allowed into the research labs at the school, halting progress and fly stock maintenance for a period. The fly stocks themselves faced some issues as well, for example the lithium concentrations used seemed to have more toxic effects than expected, causing a large portion of the flies (especially the groups with AD) to die. Along with that, the research was halted by the contamination of the UAS fly stocks, making a large portion of the UAS stocks difficult to cross, and may have led to some ineffective crosses. These incorrect crosses may also explain the outlier seen in the disease control of Table 6, contributing to the high standard error of the group. Each of these factors ultimately contributed to the low sample size and high error measurements collected from this experiment, likely contributing to the insignificant difference found from testing.

Some assumptions were also made regarding the experiment. For example, the lithium salt concentrations used were extrapolated from past research on Li_2_CO_3_ and assumed to be safe and effective for *Drosophila melanogaster* (Wang et al., 2022), and as such no toxicity assay was conducted for the Alzheimer’s flies. The lack of a toxicity assay may have contributed to the issues with fly stocks mentioned above, as the lithium dosage may be too high for the AD model used.

## Conclusion

Graphical analysis of behavioral assay performance in Figure 5 and Figure 6 suggested that AD flies treated with 5 mM LiOr performed better on average than those treated with 5 mM Li_2_CO_3_, while the reverse was true at 10 mM. These trends were consistent with the original hypothesis, but lacked statistical significance and must be interpreted with caution. However, the hypothesis that LiOr is more effective than Li_2_CO_3_ at lower dosages (5 mM), but less effective at higher dosages (10 mM), was rejected because no significant differences were observed at an alpha level of 0.05 in experimental data. Factors such as limited sample size and lack of effective control groups makes the result inconclusive, indicating that future research investigating LiOr may be warranted.

Future research should prioritize expanding sample sizes and including additional behavioral assays to elaborate on these preliminary findings. A clear extension of this experiment would be to complete data collection and get enough samples to draw statistically significant results, if possible, to get a better understanding of the effects of either lithium salts. Beyond this experiment, research could also investigate other lithium drugs (such as lithium citrate or lithium sulfite) for effectiveness in AD at higher or lower doses, specifically for clarified therapeutic windows of both lithium drugs. Alternatively, lithium drugs could be investigated in other neurodegenerative diseases (for instance Huntington’s and Parkinson’s) for similar properties or established effects in Bipolar Disorder. Beyond behavioral assays, studies could examine the molecular or cellular mechanisms of lithium action by analyzing and quantifying the Aβ and hyperphosphorylated tau levels using a western blot or ELISA, and dissecting fly brains, to get a further understanding of how the lithium salts actually affect AD.

## Supporting information

Supplementary Poster

## References

Alzheimer’s Association (2023). 2023 Alzheimer’s disease facts and figures. Alzheimer’s & Dementia, 19(4), 1598–1695. 10.1002/alz.13016

Alzheimer’s Association. (n.d.). Dementia vs. Alzheimer’s disease: What is the difference? Alzheimer’s Disease and Dementia. https://www.alz.org/alzheimers-dementia/difference-between-dementia-and-alzheimer-s

Betterlife.com. (2019). Lithium Orotate 120 mg 200 tabs, made by advanced research NCI (Dr. Hans Nieper). https://www.betterlife.com/product/lithium-orotate-120-mg/advanced-research-nci-dr-hans-nieper/62531

Bloomington Drosophila Stock Center. (n.d.). Bloomington Drosophila Stock Center Search: 4414. https://bdsc.indiana.edu/Home/Search?presearch=4414

Bloomington Drosophila Stock Center. (n.d.). Bloomington Drosophila Stock Center Search: 33774. https://bdsc.indiana.edu/Home/Search?presearch=33774

Brown, K. M., & Tracy, D. K. (2013). Lithium: The pharmacodynamic actions of the amazing ion. Therapeutic Advances in Psychopharmacology, 3(3), 163–176. 10.1177/2045125312471963

Castillo-Quan, J., Li, L., Kinghorn, K., Ivanov, D., Tain, L., Slack, C., Kerr, F., Nespital, T., Thornton, J., Hardy, J., Bjedov, I., & Partridge, L. (2016). Lithium promotes longevity through GSK3/NRF2-dependent Hormesis. Cell Reports, 15(3), 638–650. 10.1016/j.celrep.2016.03.041

Goldstein, L. E., Moir, R. D., Lee, J. C., Tabb, L. P., Saunders, A. J., & Marenda, D. R. (2011). Characterization of a *Drosophila* Alzheimer’s disease model: pharmacological rescue of cognitive defects. PLoS ONE, 6(6). 10.1371/journal.pone.0020799

Eliason, J. (2024, September 3). Drosophila Boot Camp. Lecture presented in Junior Research Class in Leesburg, Virginia.

Forlenza, O. V., De-Paula, V. J., & Diniz, B. S. (2014). Neuroprotective effects of lithium: Implications for the treatment of Alzheimer’s disease and related neurodegenerative disorders. ACS Chemical Neuroscience, 5(6), 443–450. 10.1021/cn5000309

Hardy, J., & Selkoe, D. J. (2002). The amyloid hypothesis of Alzheimer’s disease: Progress and problems on the road to therapeutics. Science, 297(5580), 353–356. 10.1126/science.1072994

Hooper, C., Killick, R., & Lovestone, S. (2007). The GSK3 hypothesis of Alzheimer’s disease. Journal of Neurochemistry, 104(6), 1433–1439. 10.1111/j.1471-4159.2007.05194.x

Kling, M. A., Manowitz, P., & Pollack, I. W. (1978). Rat brain and serum lithium concentrations after acute injections of lithium carbonate and orotate. Journal of Pharmacy and Pharmacology, 30(1), 368–370. 10.1111/j.2042-7158.1978.tb13258.x

Lee, R. M., & Koh, T. (2023). Genetic modifiers of synucleinopathies—lessons from experimental models. Oxford Open Neuroscience, 2, 1–30. 10.1093/oons/kvad001

Mirzoyan, Z., Sollazzo, M., Allocca, M., Valenza, A. M., Grifoni, D., & Bellosta, P. (2019). *Drosophila melanogaster*: A model organism to study cancer. Frontiers in Genetics, 10. 10.3389/fgene.2019.00051

National Institute on Aging. (2024). What causes Alzheimer’s disease? https://www.nia.nih.gov/health/alzheimers-causes-and-risk-factors/what-causes-alzheimers-disease

Pacholko, A. G., & Bekar, L. K. (2021). Lithium Orotate: A superior option for lithium therapy? Brain and Behavior, 11(8). 10.1002/brb3.2262

Pérez de Mendiola, X., Hidalgo-Mazzei, D., Vieta, E., & González-Pinto, A. (2021). Overview of lithium’s use: A nationwide survey. International Journal of Bipolar Disorders, 9(1). 10.1186/s40345-020-00215-z

Prüßing, K., Voigt, A., & Schulz, J. B. (2013). *Drosophila melanogaster* as a model organism for Alzheimer’s disease. Molecular Neurodegeneration, 8(1), 2–11. 10.1186/1750-1326-8-35

Rocca, W. A., Petersen, R. C., Knopman, D. S., Hebert, L. E., Evans, D. A., Hall, K. S., Gao, S., Unverzagt, F. W., Langa, K. M., Larson, E. B., & White, L. R. (2011). Trends in the incidence and prevalence of Alzheimer’s disease, dementia, and cognitive impairment in the United States. Alzheimer’s & Dementia: The Journal of the Alzheimer’s Association, 7(1), 80–93. 10.1016/j.jalz.2010.11.002

Shen, Y., Zhao, M., Zhao, P., Meng, L., Zhang, Y., Zhang, G., Taishi, Y., & Sun, L. (2024). Molecular mechanisms and therapeutic potential of lithium in Alzheimer’s disease: repurposing an old class of drugs. Frontiers in Pharmacology, 15, 1–22. 10.3389/fphar.2024.1408462

Smith, D. F., & Schou, M. (1979). Kidney function and lithium concentrations of rats given an injection of lithium orotate or lithium carbonate. Journal of Pharmacy and Pharmacology, 31(1), 161–163. 10.1111/j.2042-7158.1979.tb13461.x

Sofola-Adesakin, O., Castillo-Quan, J. I., Rallis, C., Tain, L. S., Bjedov, I., Rogers, I., Li, L., Martinez, P., Khericha, M., Cabecinha, M., Bähler, J., & Partridge, L. (2014). Lithium suppresses Aβ pathology by inhibiting translation in an adult drosophila model of Alzheimer’s disease. Frontiers in Aging Neuroscience, 6, 190. 10.3389/fnagi.2014.00190

Tan, F. H. P., Najimudin, N., Watanabe, N., Shamsuddin, S., & Azzam, G. (2023). p-Coumaric acid attenuates the effects of Aβ42 in vitro and in a *Drosophila* Alzheimer’s disease model. Behavioural Brain Research, 452, Article 114568. 10.1016/j.bbr.2023.114568

Torre, M., Bukhari, H., Nithianandam, V., Zanella, C. A., Mata, D. A., & Feany, M. B. (2023). A *Drosophila* model relevant to chemotherapy-related cognitive impairment. Scientific Reports, 13(1), Article 19290. 10.1038/s41598-023-46616-9

Tue, N. T. (2020). Insights from *Drosophila melanogaster* model of alzheimer s disease. Frontiers in Bioscience, 25(1), 134–146. 10.2741/4798

Wang, R., Ma, B., Shi, K., Wu, F., & Zhou, C. (2023). Effects of lithium on aggression in *Drosophila*. Neuropsychopharmacology, 48(5), 754–763. 10.1038/s41386-022-01475-2

