## Supplementary Poster for "Comparison of the effects of lithium orotate and lithium carbonate on locomotion and memory in a *Drosophila melanogaster* model of Alzheimer’s disease"

### Purpose

This experiment examines the difference in effects of lithium orotate and lithium carbonate as treatments for Alzheimer's disease via examining the symptoms of locomotion and memory in a *Drosophila melanogaster* model

### Background

**Alzheimer's disease (AD)**

- A **neurodegenerative** disease that affects about **1** in **9** people over the age of 65
- Symptoms include loss of **memory** and **motor function**
- These symptoms make it very difficult for AD patients to live their normal lives
- Hypothesized to be caused by the **buildup** of **Amyloid-beta (Aβ) plaques** and **tau tangles** in the nervous system
- These are irregular protein plaques that disrupt the normal functions the brain needs to conduct
- Leads to the upregulation of enzymes, causing neuroinflammation
- Neuroinflammation damages brain tissue and disrupts neural connections, accelerating AD progression
- **No definitive cure,** some treatments available to reduce symptoms

**Lithium**

- Lithium salts are currently used as treatments for bipolar disorder
- Inhibition of the GSK-3 enzyme that is upregulated during AD
- Decrease tau and Aβ accumulation, inflammation, and oxidative stress

**Lithium Carbonate ($Li_2CO_3$)**

- **Most widespread** lithium drug and the gold standard for bipolar disorder
- Has had some positive benefits on AD models in the past; however, extended usage has been shown to cause neurotoxicity

**Lithium Orotate ( LiOr)**

- **More bioavailable** than lithium carbonate meaning that it has more lithium-ion transfer to the brain
  - With equivalent dosages, LiOr was found to have 3x more ion transfer to the brain in the 1970s
- Lithium orotate 's renal toxicity at higher dosages inhibited the results
- Current studies involving lithium orotate on Alzheimer's disease are flawed and outdated

***Drosophila* as a model**

- Short life cycle and fast reproduction rate allow for large sample size
- Relatively easy gene manipulation via systems such as GAL4/UAS
  - Allows for the replication of AD
- Preserve many mammalian processes, including some neurological processes

### Importance

Alzheimer's disease is a widespread issue that can ruin the lives of millions of people worldwide. As of now, there is no cure for AD, and once diagnosed the disease is terminal and eventually fatal. Lithium salts such as lithium carbonate and lithium orotate have shown potential as an AD treatment in the past, however research on lithium orotate has been flawed and outdated If either salt is found to work as an effective treatment of AD at any concentration tested, lithium salts could pave the way to a brighter future for those diagnosed with Alzheimer's disease.

### Hypothesis

If a variety of dosages of lithium orotate and lithium carbonate are used to treat Alzheimer's disease flies, lithium orotate will be a **more effective** treatment than lithium carbonate on AD **at lower doses** (5 mM) because of the increase transfer of lithium ions to the brain, while carbonate may be better than orotate at a higher dose (10mM) due to the increased toxicity of lithium orotate.

### Experimental Method

**Independent Variables**

- Lithium drug used (lithium orotate & lithium carbonate)
- Lithium drug dosage (5 mM, 10 mM, & 20 mM)
- Presence of AD (GAL4 vs. GAL4/UAS System Cross)

**Dependent Variables**

- Locomotion (Negative Geotaxis/Climbing Assay)
- Memory (APS Assay)

**Model**

| | UAS-APP | UAS-APP |
|---|---|---|
| Act5c-GAL4 | Act5c-GAL4/ UAS-APP | Act5c-GAL4/ UAS-APP |
| CyO | CyO/ UAS-APP | CyO/ UAS-APP |

Healthy flies: GAL4
Alzheimer's flies: GAL4 x UAS

**UAS/GAL4 System Cross**

- GAL4 – Transcriptional activator that specifies location of UAS gene; Act5c expresses ubiquitously
- UAS – enhancer/expressed gene; Arctic model of Aβ-42/APP UAS expresses Aβ proteins
- Act5c-GAL4/UAS-APP progeny models Alzheimer's Disease

**Cross Steps**

1. Cross between a female UAS and a male GAL4
2. Progeny with CyO have curly wings, which was used to eliminate CyO/UAS-APP flies
3. Obtain Act5c-GAL4/UAS-APP Arctic Aβ-42 Model

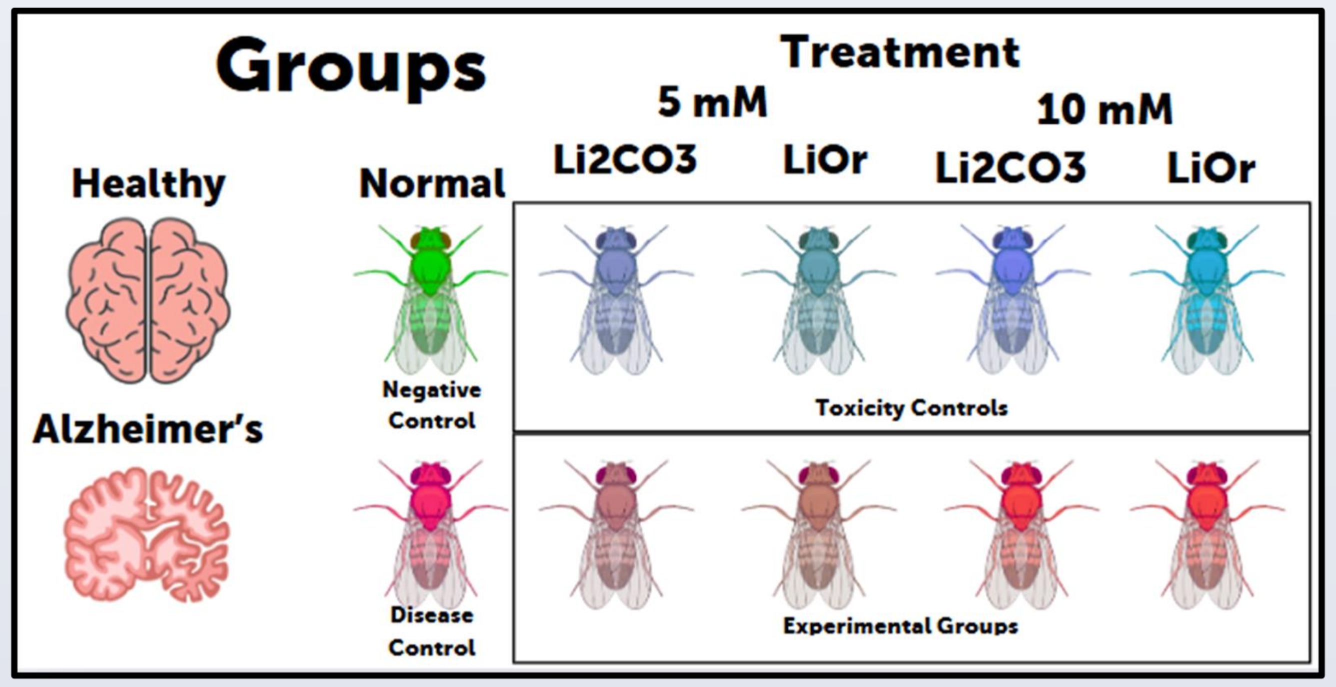


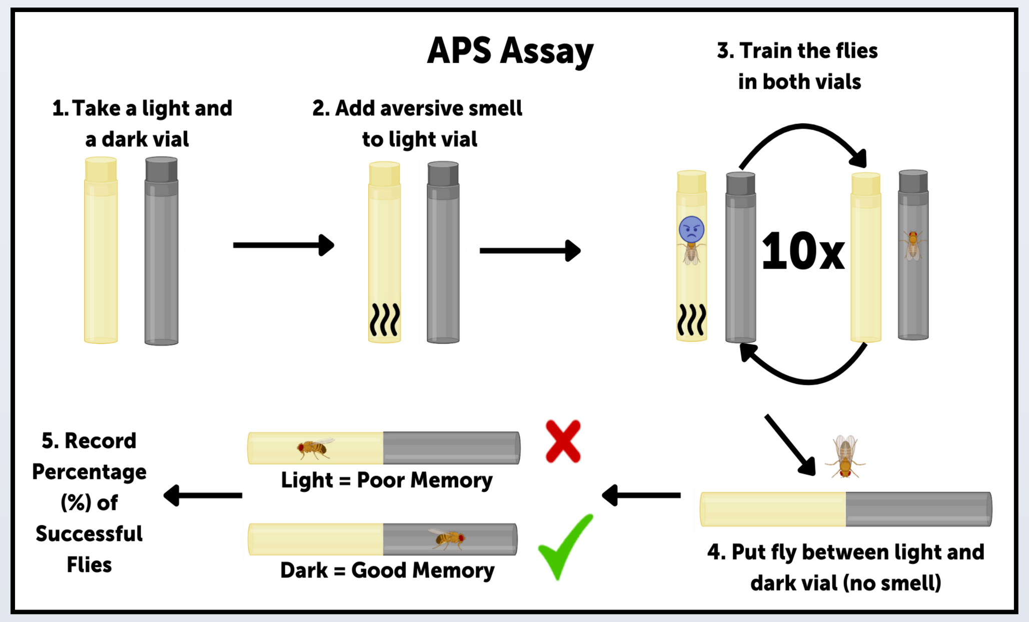


Evaluates memory based on ability to remember and avoid aversive stimulus

Negative Control
-Healthy flies in normal food
Disease Control
-AD flies in normal food
Toxicity Controls
-Varying dosages of $Li_2CO_3$ and LiOr (5mM & 10mM) in healthy flies

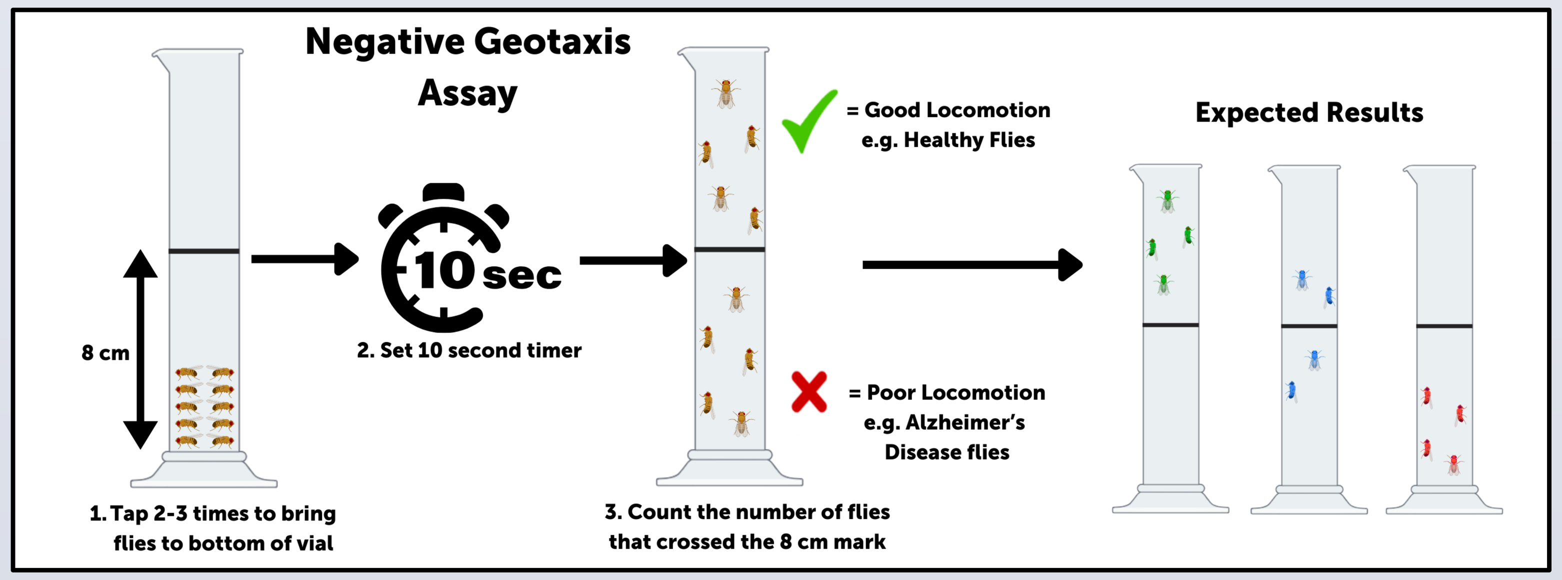


Evaluates locomotion based on ability to climb up a vial in 10 seconds

### Data

Successful flies in APS Assay (%)

| Groups | Normal Gal4 | 5mM LiOr Gal4 | 5mM $Li_2CO_3$ Gal4 | 10mM LiOr Gal4 | 10mM $Li_2CO_3$ Gal4 | Normal AD | 5mM LiOr AD | 5mM $Li_2CO_3$ AD | 10mM LiOr AD | 10mM $Li_2CO_3$ AD |
|---|---|---|---|---|---|---|---|---|---|---|
| Trial 1 | 80.00 | 72.73 | 66.67 | 81.82 | 71.43 | 12.50 | 44.44 | 40.00 | 45.45 | 55.56 |
| Trial 2 | 90.91 | 81.82 | 66.67 | 72.73 | 80.00 | 71.43 | 44.44 | 44.44 | 50.00 | N/A |
| Trial 3 | 87.50 | N/A | 63.64 | N/A | N/A | 11.11 | N/A | N/A | N/A | N/A |
| Trial 4 | 83.33 | N/A | N/A | N/A | N/A | 28.57 | N/A | N/A | N/A | N/A |
| Trial 5 | 88.89 | N/A | N/A | N/A | N/A | 22.22 | N/A | N/A | N/A | N/A |
| **Average** | **86.13** | **77.27** | **65.66** | **77.27** | **75.71** | **29.17** | **44.44** | **42.22** | **47.73** | **55.56** |

Successful flies in Climbing Assay (%)

| Groups | Normal Gal4 | 5mM LiOr Gal4 | 5mM $Li_2CO_3$ Gal4 | 10mM LiOr Gal4 | 10mM $Li_2CO_3$ Gal4 | Normal AD | 5mM LiOr AD | 5mM $Li_2CO_3$ AD | 10mM LiOr AD | 10mM $Li_2CO_3$ AD |
|---|---|---|---|---|---|---|---|---|---|---|
| Trial 1 | 71.43 | 44.44 | 70.00 | 63.64 | 66.67 | 33.33 | 40.00 | 44.44 | 44.44 | 70.00 |
| Trial 2 | 63.64 | 80.00 | 81.82 | 72.73 | 80.00 | 27.27 | 45.45 | 33.33 | 42.86 | N/A |
| Trial 3 | 80.00 | N/A | 66.67 | N/A | N/A | N/A | N/A | N/A | N/A | N/A |
| Trial 4 | 81.82 | N/A | 55.56 | N/A | N/A | N/A | N/A | N/A | N/A | N/A |
| Trial 5 | N/A | N/A | 71.43 | N/A | N/A | N/A | N/A | N/A | N/A | N/A |
| **Average** | **74.22** | **62.22** | **69.09** | **68.18** | **73.33** | **30.30** | **42.73** | **38.89** | **43.65** | **70.00** |

*Note.* Darker shades represent higher concentrations of lithium; blue represents healthy, and red represents Alzheimer's disease.

### Results

| Comparison | APS Assay *p* value | Negative Geotaxis *p* value |
|---|---|---|
| Healthy vs. AD | 0.0121 | 0.405 |
| $Li_2CO_3$ dosages | 0.0434 | 0.6121 |
| LiOr dosages | 1.0000 | 1.0000 |
| 5 mM $Li_2CO_3$ vs. LiOr | 0.1421 | 0.8631 |
| 10 mM $Li_2CO_3$ vs. LiOr | 0.6742 | 0.6742 |

| Comparison | APS Assay *p* value | Negative Geotaxis *p* value |
|---|---|---|
| $Li_2CO_3$ dosages in AD | 0.2615 | 0.1824 |
| LiOr dosages in AD | 0.2185 | 1.0000 |
| 5 mM $Li_2CO_3$ vs. LiOr in AD | 0.6251 | 0.6666 |
| 10 mM $Li_2CO_3$ vs. LiOr in AD | 0.6685 | 0.6685 |

### Results Cont.

- Healthy vs. AD & $Li_2CO_3$ vs. LiOr: p values calculated using the Mann-Whitney U test comparing APS and negative geotaxis assay pass rates between healthy and AD flies & $Li_2CO_3$ and LiOr at the given dosage
- Normal $Li_2CO_3$ or LiOr dosages: *p* values calculated using the Kruskal-Wallis test comparing APS and negative geotaxis assay pass rates between healthy flies, 5 mM lithium drug healthy flies, & 10 mM lithium drug healthy flies with the given lithium drug
- AD $Li_2CO_3$ or LiOr dosages: *p* values calculated using the Kruskal-Wallis test comparing APS and negative geotaxis assay pass rates between AD flies, 5 mM lithium drug AD flies, & 10 mM lithium drug AD flies with the given lithium drug

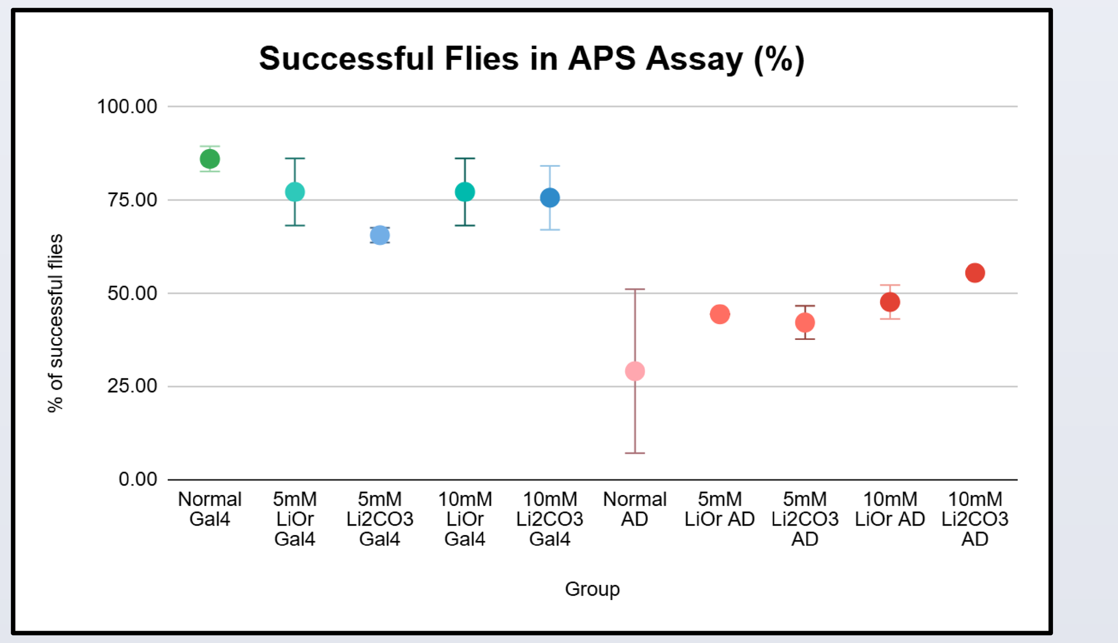


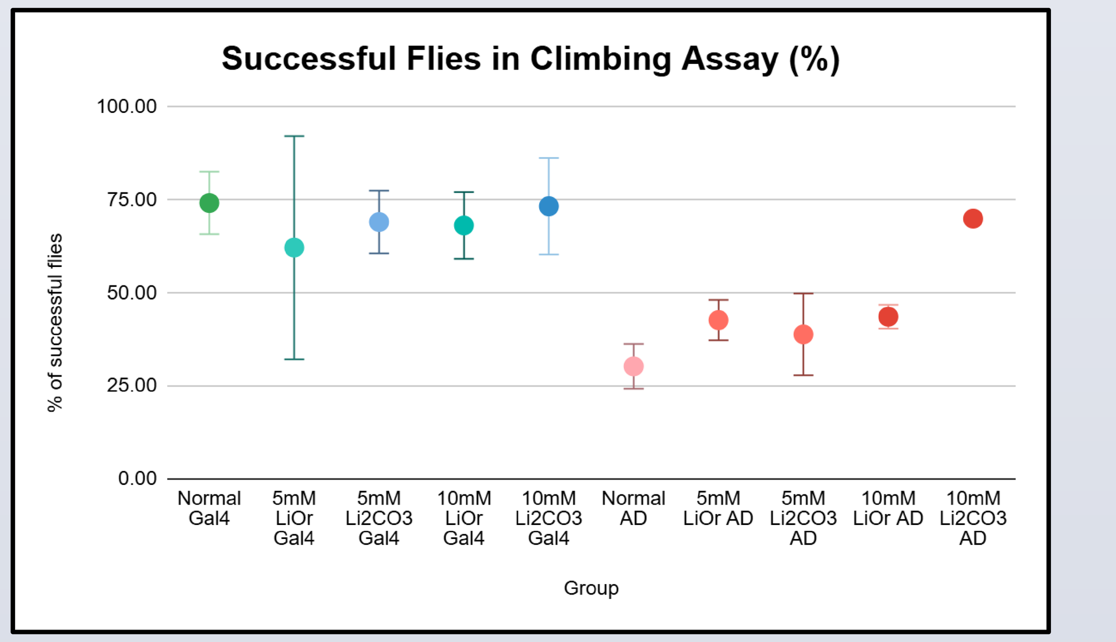


### Conclusion

- A difference was found between the memory of healthy flies and healthy flies treated with lithium carbonate, **supporting** established benefits of **lithium drugs** on **healthy** humans at **lower dosages**.
- However, lithium carbonate or lithium orotate **did not significantly** affect the locomotion of healthy flies or the locomotion or memory of AD flies.
- Ongoing data collection focuses on **completing all five trials** for the experimental and control groups
- Future research could investigate **other lithium drugs** for effectiveness in AD at higher or lower doses specifically. Alternatively, lithium drugs could be investigated in other neurodegenerative diseases for similar properties or established effects in Bipolar Disorder. Rather than looking at more subjective symptoms, looking at underlying molecular markers could yield more significant results.

### References


*Biorender*. BioRender App. (n.d.). https://app.biorender.com/

Castillo-Quan, J. I., Li, L., Kinghorn, K. J., Ivanov, D. K., Tain, L. S., Slack, C., Kerr, F., Nespital, T., Thornton, J., Hardy, J., Bjedov, I., & Partridge, L. (2016). Lithium promotes longevity through GSK3/nrf2-dependent hormesis. *Cell Reports*, *15*(3), 638–650. https://doi.org/10.1016/j.celrep.2016.03.041

Diniz, B., Machado-Vieira, R., & Forlenza. (2013). Lithium and neuroprotection: Translational evidence and implications for the treatment of neuropsychiatric disorders. *Neuropsychiatric Disease and Treatment*, 493. https://doi.org/10.2147/ndt.s33086

Forlenza, O. V., De-Paula, V. J., & Diniz, B. S. (2014). Neuroprotective effects of lithium: Implications for the treatment of alzheimer's disease and related neurodegenerative disorders. *ACS Chemical Neuroscience*, *5*(6), 443–450. https://doi.org/10.1021/cn5000309

Kocahan, S., & Doğan, Z. (2017). Mechanisms of alzheimer's disease pathogenesis and prevention: The brain, neural pathology, N-methyl-D-aspartate receptors, tau protein and other risk factors. *Clinical Psychopharmacology and Neuroscience*, *15*(1), 1–8. https://doi.org/10.9758/cpn.2017.15.1.1

Manjila, S., & Hasan, G. (2018). Flight and climbing assay for assessing motor functions in drosophila. *BIO-PROTOCOL*, *8*(5). https://doi.org/10.21769/bioprotoc.2742

*Milliporesigma*. MilliporeSigma | Life Science Products & Service Solutions. (n.d.). https://www.sigmaaldrich.com/US/en

Pacholko, A. G., & Bekar, L. K. (2021). Lithium Orotate: A superior option for lithium therapy? *Brain and Behavior*, *11*(8). https://doi.org/10.1002/brb3.2262

Pérez de Mendiola, X., Hidalgo-Mazzei, D., Vieta, E., & González-Pinto, A. (2021). Overview of lithium's use: A nationwide survey. *International Journal of Bipolar Disorders*, *9*(1). https://doi.org/10.1186/s40345-020-00215-zTue, N. T. (2020). Insights from *Drosophila melanogaster* model of alzheimer s disease. *Frontiers in Bioscience*, *25*(1), 134–146. https://doi.org/10.2741/4798


www.PosterPresentations.com
